# Vitamin D3 Regulates Mitochondrial Function and Redox Homeostasis in the Brain

**DOI:** 10.64898/2026.02.05.704094

**Authors:** Ludmila Araújo Rodrigues Lima, Pedro Lourenzo Oliveira Cunha, Heberty Tarso Facundo, Glauce Socorro de Barros Viana

**Author notes:** **Corresponding author:** Dr. Glauce Socorro de Barros Viana, Federal University of Ceará Fortaleza-Brazil.

## Abstract

Mitochondria are essential for metabolic homeostasis and neuronal function, extending beyond ATP production to roles in cell signaling, inflammation, and stress responses. Mitochondrial dysfunction, marked by abnormal morphology, ATP deficiency, and oxidative stress, is a key feature of aging-related diseases and neurodegenerative disorders like Parkinson’s. Given the importance of mitochondrial homeostasis to brain function, this study aimed to determine the possible vitamin D (VD3) effects on mitochondrial susceptibility to Ca^2+^-induced mitochondrial permeability transition pore (mPTP), bioenergetics in brain mitochondria, and redox balance. We demonstrated that VD3 protects isolated brain mitochondria. Male rats were divided into control and VD3-treated groups. Brain mitochondria were isolated for assessments of Ca^2+^-induced mitochondrial swelling secondary to MPTP opening, oxygen consumption (states 3 – ADP-stimulated and state 4 - in the presence of oligomycin), and the respiratory control ratio (RCR). Oxidative stress parameters (nitrite and lipid peroxidation), superoxide dismutase (SOD) activity, and reduced glutathione (GSH) levels were also evaluated. The results revealed that VD3 treatment blocked Ca^2+^-induced mitochondrial swelling secondary to MPTP opening. Additionally, VD3 improved mitochondrial RCR compared to controls, in the presence of complex I (malate/glutamate) and complex II (succinate) substrates, reduced mitochondrial succinate-driven H_2_O_2_ release, and enhanced SOD activity and GSH levels. These changes occurred in parallel with decreased nitrite and TBARS formation. These results suggest that vitamin D□ confers mitochondrial neuroprotection, emphasizing its prospective role in maintaining neuronal homeostasis and mitigating neurodegenerative processes.

## Introduction

Mitochondria play a central role in sustaining cellular energy balance and supporting the physiological demands of the nervous system. Through oxidative phosphorylation, they supply ATP required for maintaining electrochemical gradients and enabling efficient synaptic communication. Far beyond their traditional description as the “powerhouses” of the cell, mitochondria are now recognized as dynamic hubs that integrate metabolic regulation with pathways governing cell survival, inflammatory signaling, and adaptive stress responses [1]. Aging-related diseases are characterized by dysfunctional mitochondria which produces excessive reactive oxygen species (ROS), are abnormally distributed and produces insufficient ATP. Additionally, mitochondria can remarkably uptake and store massive amounts of calcium [2]. These factors collectively lead to subsequent damage and oxidation of proteins, lipids, and nucleic acid [3]. Moreover, impaired mitochondrial function is recognized as a key contributor to the development of several neurodegenerative diseases, notably Parkinson’s disease (PD) [4]. In the context of PD, aberrant levels of apoptotic proteins, such as alpha-synuclein, are regulated by mitochondria. lncreased ROS levels initiate a cascade culminating in non-apoptotic cell death [5].

Vitamin D3 is protective against a rat model of Parkinsonian pathogenesis induced by 6-hydroxydopamine (6-OHDA) by preserving mitochondrial function [6]. Given that mitochondrial dysfunction is a significant contributor to neurodegenerative conditions, these findings strongly suggest that VD3 could represent a promising therapeutic strategy for disorders like PD. VD3 exerts pleiotropic effects that contribute to cellular homeostasis, and epidemiological evidence links VD3 deficiency to various human diseases. Vitamin D3 (VD3) modulates autophagic activity via both genomic and rapid non-genomic signaling mechanisms. Through these pathways, VD3 exerts broad regulatory effects on diverse physiological processes across multiple organ systems, extending well beyond its classical functions in calcium homeostasis and skeletal maintenance [7].

Vitamin D3 (VD3), a biologically active secosteroid hormone is essential for maintaining physiological homeostasis and imposes protective actions in numerous organ systems. VD3 contributes to the regulation of inflammatory processes and mitigates excessive intracellular oxidative stress [8]. As a member of the nuclear steroid transcription regulator superfamily, VD3 exerts transcriptional control over numerous genes. Growing evidence indicates that VD3 may be a crucial modulator of brain development, with potential implications for neurological and neuropsychiatric disorders [9]. Furthermore, VD3 is a key regulator of systemic inflammation, oxidative stress (OS), thereby impacting the human aging process. Mitochondrial-generated adenosine triphosphate (ATP) is vital in the nervous system for establishing appropriate electrochemical gradients and ensuring reliable synaptic transmission [10].

Vitamin D3 (VD3) interacts with the vitamin D receptor (VDR), a zinc-finger transcription factor and as the other steroid hormones, VD3 is biologically active by influencing cellular physiology via both genomic and rapid non-genomic mechanisms. The genomic effects are driven by nuclear VDR-dependent transcriptional regulation, whereas the rapid signaling responses arise from VDR localized outside the nucleus. VDR expression has been identified in the brain across developmental stages as well as in adulthood, supporting its essential role in modulating neurodevelopmental processes and maintaining neural function [11]. VDR is fundamental for many of the vitamin’s genomic and non-genomic effects, and the absence of this receptor compromises mitochondrial integrity and leads to cell death, highlighting the pivotal role of the VDR in safeguarding cells against excessive ROS generation and the consequent oxidative damage [12]. Therefore, the objectives of the present study were to investigate the effects of VD3 on isolated brain mitochondrial function from rats subchronically treated with VD3. The primary focus was on assessing Ca^2+^-induced mitochondrial swelling, respiration, and oxidative stress.

## Materials and methods

### Animals and Experimental design

Adult male Wistar rats (∼250 g) were maintained under controlled environmental conditions on a 12 h light/dark cycle, with unrestricted access to standard laboratory chow and water (Ad libitum). Animals were randomly assigned to experimental groups, 5 to 6 rats per group: 1. Control group and 2. VD3 group. The control group received daily oral gavage administration of 0.2 mL 1% DMSO in distilled water. Animals in the VD3 group were administered Vitamin D (VD3 - Sanofi, São Paulo) via oral gavage at a dose of 1 μg/kg/day for 14 consecutive days, with administrations consistently performed at 10:00 AM. The VD3 was suspended in a 1% aqueous DMSO solution prior to administration. The selected VD3 dosage was based on previous studies demonstrating its neuroprotective efficacy [13]. Following the treatment period, all animals were anesthetized using 100 mg/kg ketamine plus 10 mg/kg xylazine prior to euthanasia.

### Mitochondrial Isolation

Brain mitochondria were isolated by following the detailed protocol described by [14]. This procedure employs differential centrifugation and yields a preparation containing both synaptosomal and non-synaptosomal mitochondria. Rat brain tissue was initially washed at 4 °C in an isolation buffer containing in mM: 10 HEPES, 10 EGTA, 125 sucrose, 250 mannitol (pH 7.2, adjusted with KOH) plus 0.01% BSA. The material was cut in small blocks and homogenized in a glass tissue grinder. The resulting homogenate was subjected to an initial centrifugation at 2,000 x g for 3 minutes to pellet intact cells and nuclei. Then, the supernatant was centrifuged twice at 12,000 x g for 8 minutes. 0.1% digitonin was added to the buffer before the second centrifugation. The resulting mitochondrial pellet was then resuspended in the isolation buffer, this time excluding EGTA. The final supernatant from this step was collected separately to serve as the cytosolic fraction for the analysis of cytosolic superoxide dismutase (SOD) activity. Sample protein concentration was quantified spectrophotometrically using the Bradford method [15], with bovine serum albumin (BSA) serving as the standard.

### Mitochondrial Swelling

Mitochondrial swelling, indicative of Ca^2^□ uptake and mitochondrial permeability transition pore (mPTP), was assessed by monitoring changes in light scattering at 540 nm. Isolated rat brain mitochondria were suspended in an assay buffer containing in mM: 10 succinate, 150 10 HEPES, KCl, 2 MgCl□, 2 KH□PO□, adjusted to pH 7.2 with KOH, plus and 1 μg/mL oligomycin (condition with high membrane potential – State 4). The mPTP was triggered by the addition of 2 μM Ca^2^□, and the resulting decline in absorbance was continuously recorded for 5 minutes.

### Mitochondrial oxygen consumption

Mitochondrial oxygen consumption was assessed using a Clark-type oxygen electrode (Oxygraph, Hansatech Instruments, UK) interfaced with a computer-based data acquisition system. The respiration buffer was composed of (in mM): 25 sucrose, 75 mannitol, 5 KH□PO□, 100 KCl, 20 Tris–HCl, pH 7.2. 1 mg/mL fatty acid–free bovine serum albumin was added to the buffer just before use. Oxidative activity was initiated with glutamate (5 mM) plus malate (2.5 mM) or succinate (10 mM) as substrates to drive complex I– or complex II–linked respiration, respectively. State 3 respiration was stimulated by the addition of ADP (500 μM), followed by induction of state 4 through the addition of oligomycin (1 μg/mL).

### Mitochondrial respiratory control ratio (RCR)

Mitochondrial efficiency, often expressed as the capacity to couple oxygen consumption to ATP synthesis, was evaluated through the respiratory control ratio (RCR). The RCR was determined as the ratio of oxygen consumption during state 3 (ADP-stimulated respiration) to that observed during state 4 (oligomycin-induced respiration). Respiration rates were calculated from the linear slope representing the negative time derivative of oxygen concentration. This parameter reflects the balance between oxidative phosphorylation activity and proton leak across the inner mitochondrial membrane.

### Measurements of mitochondrial H_2_O_2_ production

Mitochondria were incubated for 30 mins with 5 µM Amplex red (Molecular Probes, Eugene, Oregon, USA) and 1 U/ml HRP to measure H_2_O_2_. The reaction was conducted in respiration buffer containing in mM: 100 KCl, 20 Tris–HCl, 25 sucrose, 75 mannitol, 5 KH_2_PO_4_, plus 1 mg/ml fatty acid-free BSA, pH 7.2. Importantly, to induce state 4 we added 1 µg/ml oligomycin. After 30 mins in 37°C the tube was centrifuged (at 5000g for 2 minutes) and the absorbance of the supernatant was read at 560 nm. The blank tube (incubated for the same conditions as the test tubes) contained Amplex red and horseradish peroxidase without the samples. Mitochondrial H_2_O_2_ production was calculated from a standard curve generated from known concentrations of H_2_O_2_.

### Determination of nitrite contents

Nitrite concentrations were measured using the Griess colorimetric assay. Briefly, tissue homogenates (10% w/v in KCl buffer) were centrifuged at 10,000 × g for 10 min. Then, the supernatant was mixed with freshly prepared Griess reagent (1 part 0.1% naphthylethylenediamine dihydrochloride in distilled water and 1 part 1% sulfanilamide in 5% H□PO□). After incubation at room temperature for 10 min, absorbance was recorded at 520 nm using a spectrophotometer. Nitrite concentrations were calculated from a sodium nitrite standard curve and expressed as micromoles of nitrite per gram of tissue [19].

### Thiobarbituric Acid Reactive Substances Detection (TBARS)

Lipid peroxidation is largely used to quantify oxidative stress [20][23]. In this study we quantified Malondialdehyde as a final product of lipid peroxidation. Briefly, homogenates were added to 10% TCA followed by the addition 0.6% thiobarbituric acid. Following vigorous mixing, the samples were incubated in a water bath at 95–100 °C for 15 min to allow color development. Subsequently, they were rapidly cooled on ice and centrifuged at 1,500 × g for 5 min. The TBARS content was quantified spectrophotometrically at 540 nm using a microplate reader, and the results were normalized as micromoles of malondialdehyde (MDA) per gram of tissue. For quantification, a standard calibration curve was constructed using known MDA concentrations.

### Determination of the concentration of reduced Glutathione (GSH)

Free thiol group from GSH was detected by using the Ellman method, using 5,5′-dithiobis (2-nitrobenzoic acid) (DTNB). This reaction was assessed using an ELISA plate reader. Briefly, the tissue homogenate (10% in phosphate buffer) was mixed with trichloroacetic acid (TCA) to precipitate proteins. The resulting mixture was centrifuged at 3,000 rpm for 15 minutes at 4°C. For the assay, the supernatant or buffer (blanc)were transferred to a cooled ELISA plate. Prior to reading at 412 nm, 0.01 M DTNB in methanol were added to each well. The final GSH concentration was derived from a standard GSH curve and is reported as μg per gram of tissue.

### Superoxide dismutase activity (SOD)

Superoxide dismutase (SOD) activity was quantified using the nitro blue tetrazolium (NBT) photoreduction assay. In brief, the reaction was conducted in potassium phosphate buffer containing in mM: 13 L-methionine, 75 NBT, and 0.1 EDTA (pH 7.8). 2 μM riboflavin was added to initiate the reaction followed by exposure for 10 minutes to uniform, unfiltered white light. The developed blue color was read at 560 nm. SOD activity was expressed as units (U) normalized per milligram of sample protein.

### Statistical analyses

All data were expressed as mean ± SEM. For all parameters, the unpaired t-test was used. p < 0.05 was considered statistically significant. All statistical analyses were performed with the GraphPad Prism (Version 7.0, USA).

## Results

### VD3 blocks Ca^2+^-induced mPTP

Mitochondrial swelling secondary to mPTP opening is one of the most critical marks of mitochondrial ultrastructural changes. mPTP opening (due to supraphysiological Ca^2+^ accumulation associated with oxidative stress) promotes depolarization, oxidative phosphorylation disturbances, swelling, and, ultimately, release cytochrome c and other apoptogenic proteins due to outer membrane rupture [21, 22]. Mitochondrial swelling was performed in suspensions of isolated mitochondria in each condition by light scattering at 540 nm. The decline in absorbance indicates an intensification in mitochondrial swelling. Mitochondria isolated from VD3 treated samples in the presence of succinate as substrate exhibited a significantly lower Ca^2+^-induced swelling than controls as shown in the representative traces of Figure 1A and averages ± standard errors in Figure 1B.

**Fig. 1.**
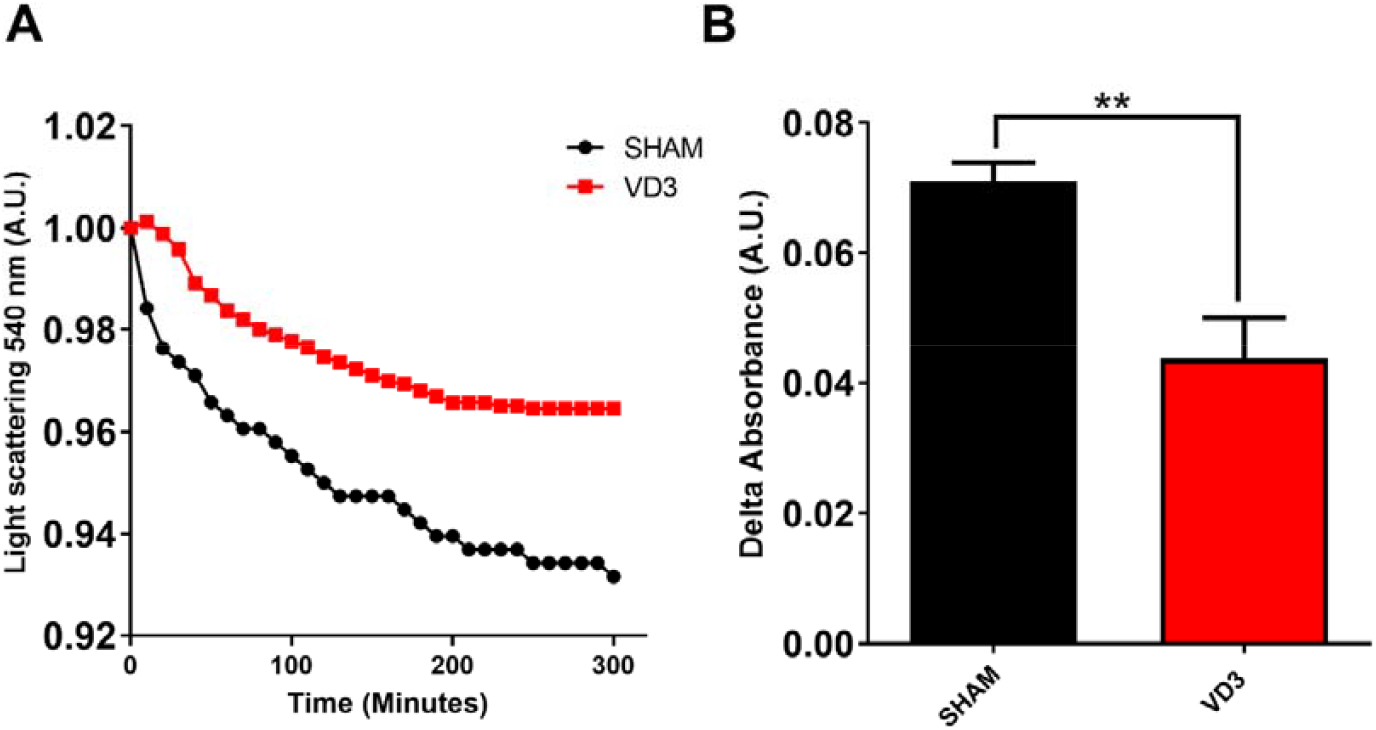
Vitamin D (VD3) attenuates calcium-induced swelling of rat brain mitochondria. 3Just before any swelling curve, 2 μM Ca^2+^ was added to induce the opening of mPTP. A. Representative traces of Ca^2+^ induced mitochondrial swelling at 540 nm for 300s. B. Delta of absorbance between 300s and 0 time point. The data are presented as mean values ± SEM (n=5). **p<0.01 (Unpaired t-test).

### VD3 positively impacts mitochondrial respiration

Oxygen consumption is a critical parameter for assessing mitochondrial respiratory activity. Based on this, we investigated whether VD3 enhances mitochondrial oxygen consumption. Our initial findings revealed that VD3 did not affect maximal respiration (state 3 respiration) induced by ADP in the presence of succinate as a substrate (Figure 2A). However, VD3 significantly enhanced state 3 respiration when malate/glutamate was used as substrates (Figure 2A). In contrast, when ATP synthase was inhibited by oligomycin (state 4 respiration), mitochondria from VD3-treated samples exhibited significantly lower respiration with succinate as the substrate (Figure 2A), while no change was observed in malate/glutamate-supported respiration. These seemingly opposing effects led to alterations in respiratory control ratios (RCRs) in both conditions (Figures 2A, and B), which reflect the coupling efficiency between respiration and ATP synthesis in mitochondria. These findings suggest that VD3 modulates mitochondrial respiratory efficiency in a distinct substrate-dependent manner, enhancing state 3 respiration with malate/glutamate and reducing state 4 respiration with succinate.

**Fig. 2.**
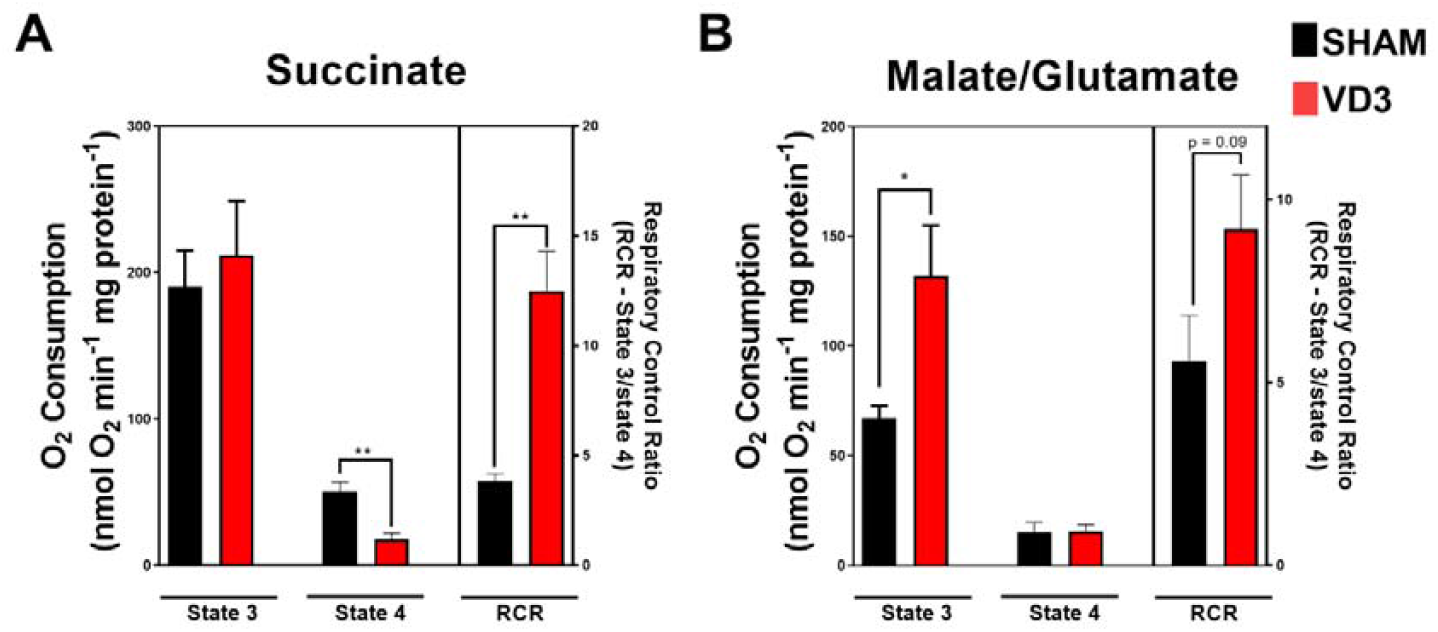
VD3 modulates brain mitochondrial oxygen consumption. Oxygen consumption rates were measured in brain isolated mitochondria incubated, successively, in the presence of substrates of complex II succinate plus ADP (A) or complex I substrate malate/glutamate plus ADP (B - State 3), and oligomycin (State 4). B. RCR = Respiratory control ratios calculated as State 3/State 4. Each bar represents mean values ± SEM (5 animals per group). *p < 0.05, **p < 0.01 Unpaired t-test.

### VD3 protects brain cells against oxidative stress

The brain is extremely sensitive to oxidative stress considering that neurons have high metabolic activity associated with a small concentration of endogenous antioxidants [23]. Accumulation of ROS causes severe damage to cellular components and is related to the etiopathogenesis of numerous neurodegenerative diseases. In mammals, mitochondria are a primary source of ROS [24, 25]. In order to test the effects of VD3 on mitochondrial oxidative stress we measured the H_2_O_2_ produced by brain isolated mitochondria in the presence of succinate plus oligomycin. It is important to note that this condition will generate high levels of mitochondrial ROS by reverse electron transport [14, 26]. Strikingly, animals that were supplemented with VD3 showed a significant reduction in mitochondrial H_2_O_2_ production (in state 4) when compared with control animals (Figure 2A).

Having seen the effects of VD3 mitochondrial H_2_O_2_ production, we tested whether this procedure had any effect on SOD activity. Interestingly, low SOD activity have been associated with several neurodegenerative diseases like alzheimer’s disease, parkinson’s disease, huntington’s disease, and amyotrophic lateral sclerosis [27]. For the determination of cytosolic SOD activity, we used NBT assay (as described in materials and methods). Strikingly, VD3 suplementation significantly enhanced cytosolic SOD activity in comparison with control group (Figure 3B).

**Fig. 3.**
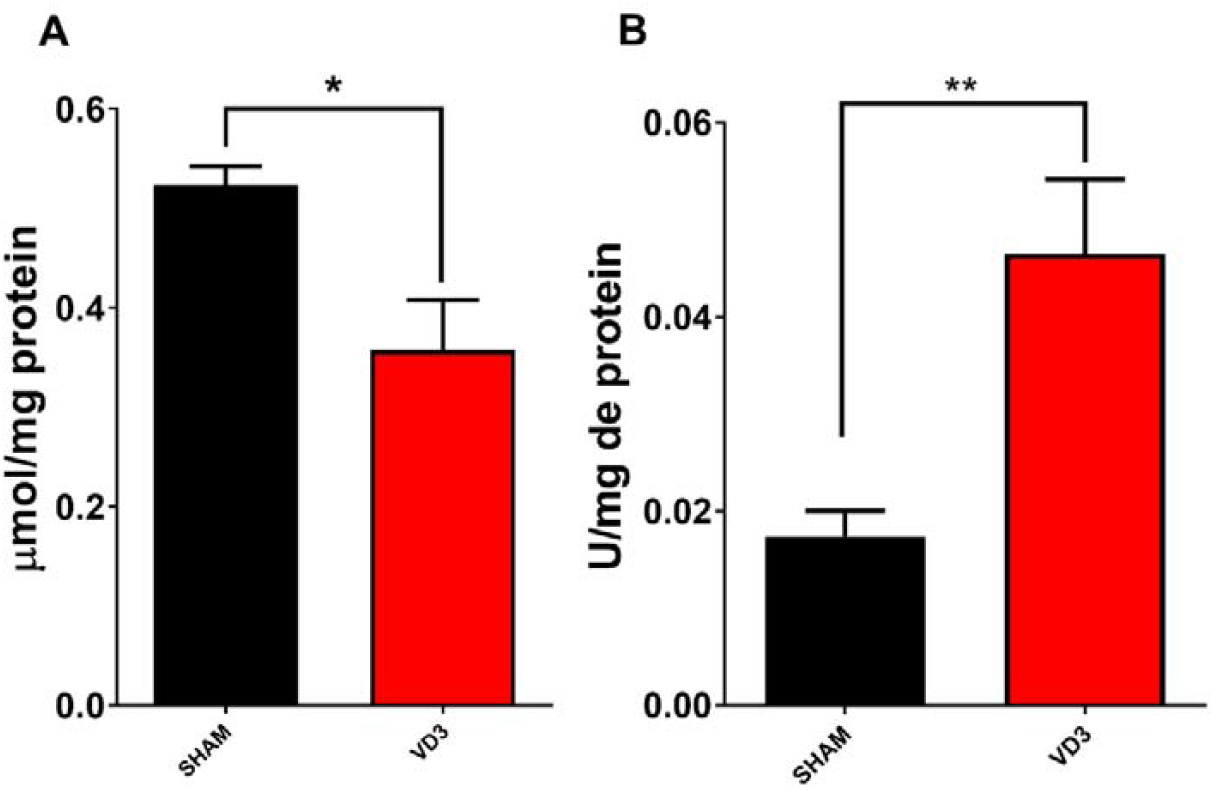
Vitamin D (VD3) blocks mitochondrial succinate-induced H_2_O_2_ release and improves superoxide dismutase (SOD) activity in brain mitochondria. A. Mitochondrial H_2_O_2_ production. B. Superoxide dismutase activity (SOD) levels. Each bar represents mean values ± SEM (5 animals per group). *p < 0.05, **p < 0.01 Unpaired t-test.

Oxidative stress which is characterized by high oxidant and low antioxidant levels triggers a series of reactions that will finally oxidize macromolecules such as lipids, proteins and DNA [28]. Knowing that, we wondered if low mitochondrial ROS production with improved SOD activity in VD3 would impact the nitrite contents, lipid peroxidation (TBARS) and GSH concentration in prefrontal cortex, hippocampus and striatum. As expected, VD3 supplementation was able of decrease nitrite (Figure 4A–D) and TBARS levels (Figure 5A – D) in all brain areas studied. VD3 also improved GSH levels (Figure 6A – D). These results indicate that VD3 supplementation protect cells against oxidative stress by improving mitochondrial function and preserving SOD activity and GSH levels. Altogether, this enhanced brain redox capacity prepare this organ against any pathological redox or mitochondrial disturbances.

**Fig. 4.**
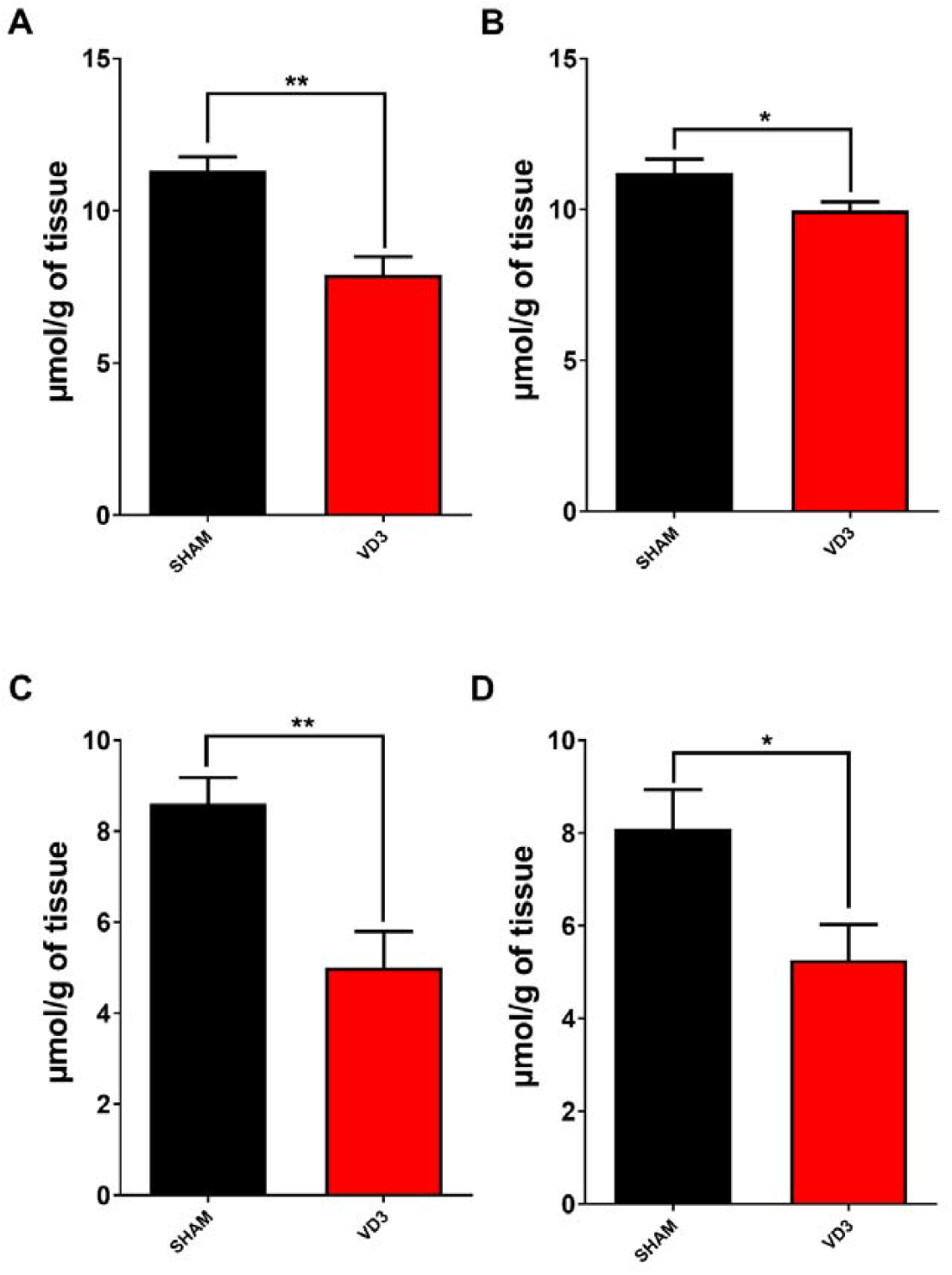
Nitrite levels in prefrontal cortex (A), hippocampus (B), right striatum (C) and left striatum (D) in controls (SHAM) and vitamin D (VD3)-treated groups. Each bar represents mean values ± SEM (5 animals per group). *p < 0.05, **p < 0.01 Unpaired t-test.

**Fig. 5.**
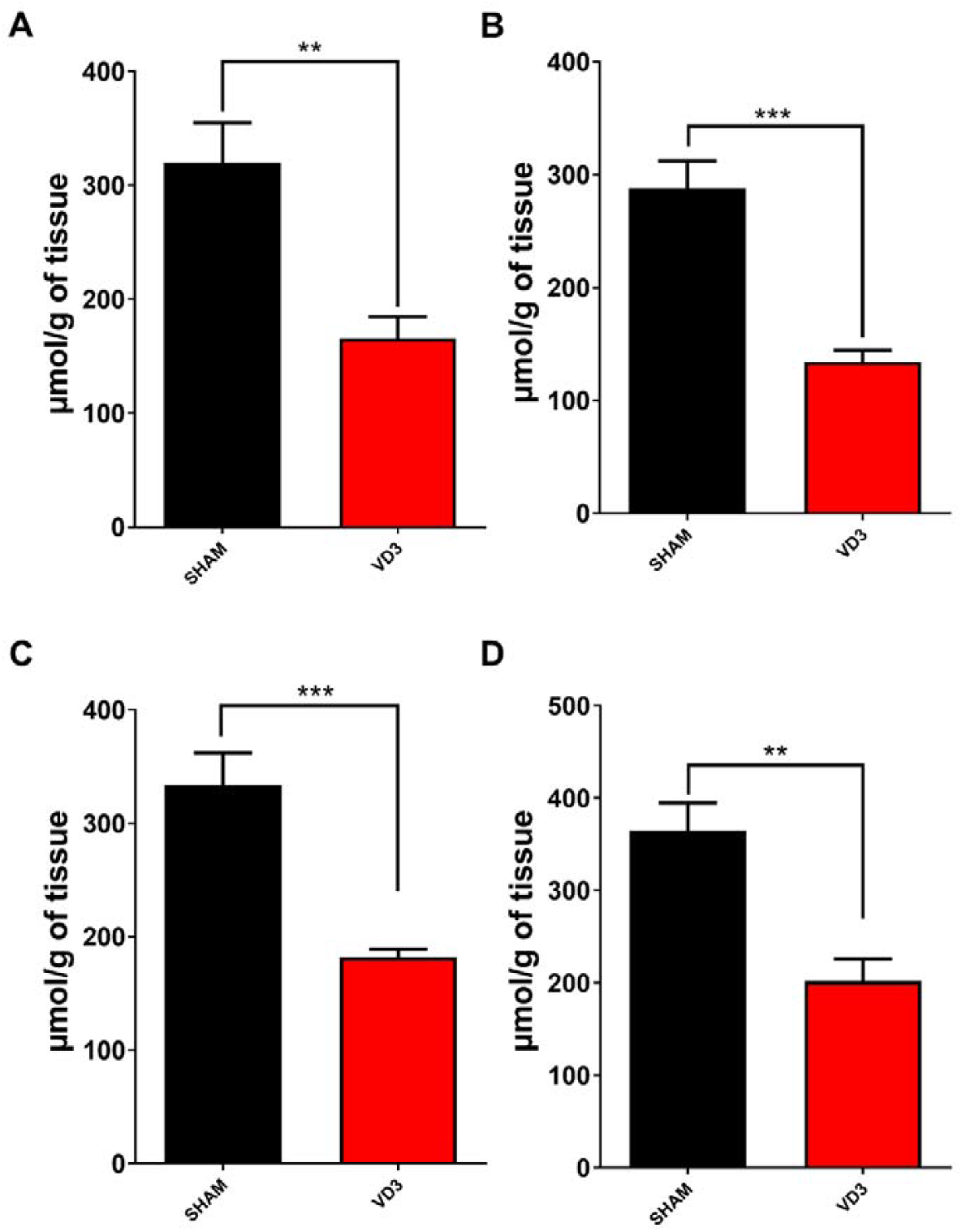
TBARS levels in prefrontal cortex (A), hippocampus (B), right striatum (C), and left striatum (D) in controls (SHAM) and vitamin D (VD3)-treated groups. Each bar represents mean values ± SEM (5 animals per group). *p< 0.05, ** p<0.01, ***p< 0.001 unpaired t-test.

**Fig. 6.**
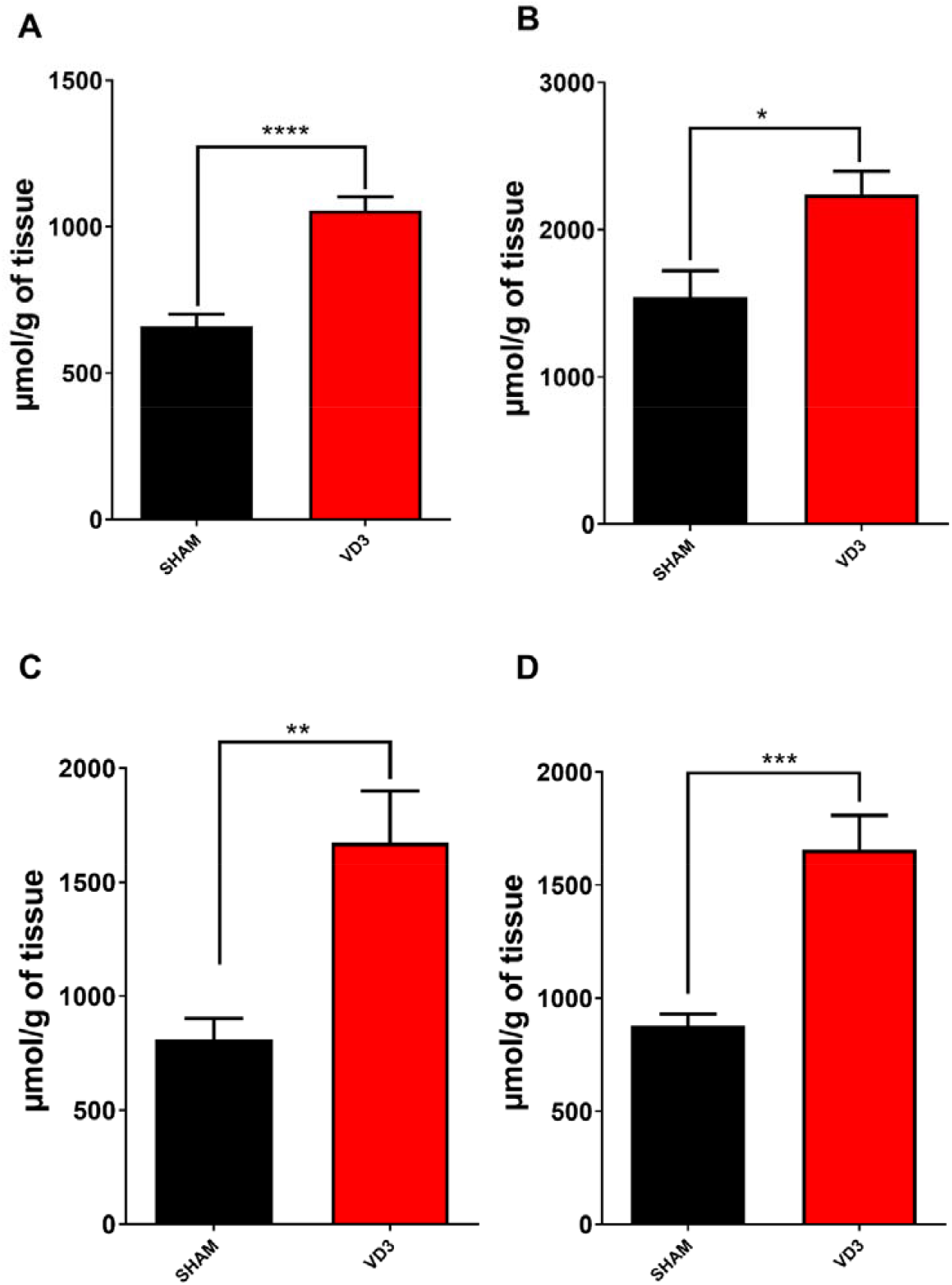
GSH levels in prefrontal cortex (A), hippocampus (B) right striatum (C), and left striatum (D) with and without vitamin D (VD3) treatment. The data are presented as mean values ± SD (5 animals per group). *p< 0.05, ** p<0.01, ***p< 0.001 Unpaired t-test.

## Discussion

The rising global life expectancy has led to a growing aging population and an increasing prevalence of neurodegenerative diseases, such as Parkinson’s disease (PD). Evidence indicates that mitochondria play a central role in these aging-related disorders, acting as key regulators of cell death. Notably, mitochondrial DNA mutations and oxidative stress contribute to aging, the primary risk factor for neurodegenerative diseases. Therefore, therapies targeting core mitochondrial processes hold significant promise [28]. Recently, we have demonstrated that vitamin D□ (VD□) exerts neuroprotective effects in brain mitochondria from hemiparkinsonian rats by reducing mitochondrial swelling and oxidative stress markers [6]. Mitochondrial dysfunction has been linked to the cytotoxic effects of 6-hydroxydopamine, a neurotoxin commonly used in PD models, which increases mitochondrial membrane permeability and triggers cytochrome c release [29, 30]. The impact of Vitamin D on mitochondrial function and oxidative stress in brain tissue under physiological conditions remains uncertain. In this study, we observed that VD3 treatment prevented Ca^2^□-induced mitochondrial swelling associated with MPTP opening. Furthermore, VD3 enhanced mitochondrial efficiency, as indicated by improved RCR with both complex I (malate/glutamate) and complex II (succinate) substrates. It also reduced succinate-driven H□O□ release, increased SOD activity and GSH levels, and lowered nitrite and TBARS formation.

Mitochondrial dysfunction is now recognized as a pivotal factor in the pathogenesis of Parkinson’s disease [31]. Mitochondria serve as central regulators of eukaryotic cell fate, governing both survival and death through oxidative phosphorylation–driven energy metabolism. Mitochondrial dysfunction, whether functional or structural, has been strongly linked to the cytotoxic effects of 6-OHDA (a synthetic neurotoxin which experimentally induces Parkinson’s disease). In Parkinson’s disease, patients exhibit deficiencies in the oxidative phosphorylation system [32]. In the present study, we demonstrated lower susceptibility of brain mitochondria to Ca^2+^-induced swelling in animals supplemented with VD3 in comparison with the control group. mPTP is a phenomenon induced by Ca^2+^ and oxidative stress that unselectively promotes the passage of small solutes through the inner mitochondrial membrane [22]. Unlike the permeabilization of the outer mitochondrial membrane, which primarily promotes apoptotic signaling, the opening of the mPTP can elicit a wide spectrum of cellular outcomes. Depending on the context, mPTP activation may contribute to normal mitochondrial quality control through mitophagy or, under pathological conditions, lead to apoptotic or necrotic cell death [33]. Mitochondrial swelling—defined as the swelling of the organelle’s matrix—arises from increased permeability of the inner membrane. This matrix expansion can exert mechanical stress on the outer membrane, ultimately causing its rupture [16]. By reducing ROS formation and increasing antioxidant buffering capacity, VD3 likely lowers the probability of mPTP opening, thereby protecting mitochondrial function under stress conditions [22]. Our results in VD3 treatment are consistent with this idea since VD3 treatment not only blocks mitochondrial ROS formation but also enhances cellular antioxidant at physiological levels.

The efficiency of mitochondrial respiration reflects the tight coupling between oxidation and phosphorylation. In this context, we observed that vitamin D□ (VD□) supplementation did not alter oxygen consumption in succinate respiring conditions (State 3). However, VD□ significantly decreased state 4 respiration under the same conditions. Conversely, when mitochondria were energized with malate plus glutamate (which supply NADH to complex I), a significant increase in oxygen consumption was observed in state 3, but not in state 4. These findings suggest that VD□ differently modulates mitochondrial respiration depending on the substrate fueling the electron transport chain. This substrate-dependent effect may arise from the distinct redox environments generated by complex I– and complex II–linked respiration. Under succinate-supported conditions, particularly in state 4, reverse electron transport (from complex II to complex I) is known to promote elevated reactive oxygen species (ROS) formation [14, 26]. VD3 might block this high ROS formation as seen in Figure 2 positively regulating mitochondrial respiration and avoiding detrimental effects of ROS in mitochondrial oxygen consumption. ROS are also formed by Complex I especially in conditions of inhibition of this complex using rotenone (ROT) [33]. A recent study [34], VD3 pretreatment was capable of preventing loss of viability in cells exposed to ROT by blocking both necrosis and apoptosis. Furthermore, cells exposed to ROT (in the absence of VD3) showed increased ROS production, in a manner blocked by VD3 treatment. Additionally, VD3 also prevented ROT-induced mitochondrial membrane potential decrease. Moreover, VD3 treatment safeguarded astrocytes against ROT-induced injury by mitigating oxidative stress and enhancing mitochondrial performance. The improvement in state 4 respiration observed with VD□ could therefore reflect a protective mechanism that limits RET-induced ROS production, as illustrated in Figure 2, thereby maintaining mitochondrial efficiency and preventing oxidative damage. This suggests that changes in H_2_O_2_ release promoted by VD3 in brain mitochondria may be a consequence of improved susceptibility to ROS at physiological basal levels. Such an effect could have pathophysiological relevance, particularly during ischemia–reperfusion events in the brain, where succinate accumulation drives excessive ROS generation upon reperfusion [35]. Thus, VD□ may confer protection by improving mitochondrial susceptibility to ROS and by modulating respiration in a substrate-dependent manner.

The mitochondrial electron transport chain drives ATP synthesis through oxidative phosphorylation, a process fundamental to cellular energy homeostasis and redox signaling [36]. Mitochondria also play a central role in apoptosis, a key pathological feature of neurodegenerative diseases such as PD, in which abnormal accumulation of the pro-apoptotic protein α-synuclein is observed. In addition, mitochondria are major sources of ROS, which, when produced in excess, can damage DNA, proteins, and lipids [37]. Besides, ROS will contribute to inflammatory pathways, excitotoxicity, protein agglomeration, and apoptosis [38]. In the present study, VD□ supplementation significantly SOD activity across brain regions. Conversely, control animals exhibited higher nitrite levels and lipid peroxidation compared with VD□-treated rats. Moreover, all brain regions from VD□-supplemented animals showed a marked elevation in reduced glutathione (GSH) content, supporting the hypothesis that VD□ exerts neuroprotective effects by enhancing the antioxidant defense system. Based on these findings, we propose that VD3 prevents oxidative stress not only by fine-tuning mitochondrial respiration/function but also by maintaining SOD and GSH levels. This interpretation is consistent with previous studies. For instance, a study reported that VD□ pretreatment prevented the decline in GSH levels in rat brain tissue from an Alzheimer’s disease (AD) model [39]. Similarly, another study [40] demonstrated that VD□ supplementation attenuated lipid peroxidation in the brains of female rats with AD, a neurodegenerative condition that shares mechanistic features with PD. Supporting evidence from cell culture studies also indicates that VD□ enhances GSH synthesis and reduces ROS and pro-inflammatory cytokine production in monocytes [41].

We recently [38] showed that physical exercise combined with VD3 treatment in 6-OHDA-lesioned hemiparkinsonian rats improved behavioral outcomes, accompanied by comparable increases in dopamine and 3,4-dihydroxyphenylacetic acid levels. Additionally, changes in tyrosine hydroxylase, dopamine transporter, and VD3 receptor immunoexpressions were minimized by VD3 plus exercise. VD□ has been shown to modulate mitochondrial redox balance, attenuating ROS generation and enhancing oxidative phosphorylation capacity [42]. Consistent with this, VD□ receptor (VDR) knockdown reduces mitochondrial respiration and ATP synthesis, underscoring its role in maintaining mitochondrial bioenergetics. Consistent with these observations, VD□ deficiency correlates with increased susceptibility to inflammatory and oxidative stress– related disorders [8, 43]. VD□ signaling involves both genomic and rapid non-genomic pathways mediated by VDR, which is expressed in the developing and adult brain [7, 9, 11, 43]. Within the CNS, mitochondrial ATP production supports synaptic transmission, while dysfunction in mitochondrial dynamics, ion homeostasis, or mitophagy contributes to neurodegenerative processes, such as PD [4, 44]. In addition, VD3 protects against cognitive decline and dementia, and higher brain VD3 concentrations were associated with better cognitive function prior to death. The interplay between mitochondrial impairment and neuroinflammatory signaling represents a multifaceted and highly dynamic process and mitochondrial dysfunction seems to precede neuroinflammation in the progression of the neurodegenerative disease [45].

## Conclusion

Beyond energy production, mitochondria integrate metabolic and epigenetic signals regulating neuronal survival. Their dysfunction precedes and amplifies neuroinflammatory cascades, reinforcing their central role in neurodegeneration [46–48]. In conclusion, our data indicate that VD□ preserves mitochondrial function—reducing swelling, enhancing respiratory control, and limiting oxidative stress—thereby mitigating key pathological features associated with neurodegenerative diseases such as PD.

## Author Contributions

All authors contributed equally to this work.

## Data Availability

All data are available on reasonable request with the corresponding author.

## Declarations

### Ethical Approval

The experimental protocol received approval from the Institutional Committee for Animal Experimentation of the Federal University of Ceara (UFC), identified by protocol number 3397010219. Furthermore, the study strictly adhered to the guidelines outlined in the 8th Edition of the Guide for the Care and Use of Laboratory Animals, published by the National Research Council, USA.

